# Optimization of a Subcutaneous Epithelioid Hemangioendothelioma Xenograft Model for Evaluation of Preclinical Therapeutics

**DOI:** 10.64898/2026.11.03.764004

**Authors:** Zilu Chen, Lucy Wang, Valentina Zuco, Silvia Stacchiotti, Sandro Pasquali, Nadia Zaffaroni, Albiruni R. Abdul Razak, Christine Allen

## Abstract

Epithelioid hemangioendothelioma (EHE) is an ultra-rare vascular sarcoma characterized by an unpredictable clinical trajectory and lack of established standard-of-care treatment regimens. Despite the pressing need for effective therapeutic strategies, the scarcity of clinically relevant preclinical models remains a major barrier to drug development. To address this challenge, we used a patient-derived human EHE cell line to optimize and validate a subcutaneous xenograft model in mice.

Optimization of cell-inoculation density revealed that injection of high cell densities was associated with adverse systemic responses and primary tumour growth kinetics unsuitable for longitudinal drug screening. In contrast, inoculation of low, sub-million cell densities generated reproducible xenografts that followed classical sigmoidal growth kinetics, providing a suitable therapeutic window for comparative treatment studies.

Subsequent histopathological analysis confirmed that the tumours were highly proliferative and retained key features of EHE pathology, including expression of the EHE-associated markers ERG and CAMTA1. Treatment response validation further demonstrated sensitivity to mTOR inhibition, a characteristic preserved from the original patient-derived sample. Treatment with nab-sirolimus significantly suppressed tumour growth and reduced phosphorylation of ribosomal protein S6, consistent with suppression of downstream mTORC1 signalling.

In this study, the optimized CDX model retains key EHE features and provides an extended observation window with sustained growth of untreated controls, supporting longitudinal assessment of antitumour activity and pharmacodynamic responses. This platform provides a practical framework for evaluating and comparing candidate therapies for EHE.

## 1. Introduction

Epithelioid hemangioendothelioma (EHE) is an ultra-rare vascular sarcoma of endothelial origin characterized by highly variable clinical behaviour and limited treatment options [1,2]. EHE may present as localized, multifocal, or metastatic disease, and its long-term clinical course ranges from indolent disease to rapid progression [3–6]. Histologically, EHE is characterized by epithelioid endothelial cells embedded within a myxohyaline stroma and is molecularly defined in most cases by the *WWTR1::CAMTA1* fusion, with a smaller subset harbouring *YAP1::TFE3* rearrangements [7–9].

Owing to the rarity and heterogeneous clinical behaviour of EHE, no systemic therapy has been established as a universally accepted standard-of-care for advanced disease. Although surgical resection may be effective for localized disease, unresectable or metastatic EHE is often managed using empiric approaches adapted from other sarcoma subtypes such as cytotoxic chemotherapy, anti-angiogenic agents, and molecular targeted therapies such as mTOR inhibitors [1,10]. Among these, the mTOR inhibitor sirolimus has demonstrated clinical activity in patients with advanced EHE, particularly in those with progressive disease without serosal effusion [11]. Nevertheless, therapeutic responses are variable, and the biological mechanisms underlying treatment sensitivity remain poorly understood.

Despite substantial effort, progress in the development of effective therapies for EHE has been hindered by the limited availability of robust preclinical models. Due to its rarity, only a small number of EHE cell lines and experimental systems have been reported. Existing studies have largely relied on engineered endothelial, fibroblast, or other non-EHE cell models expressing *WWTR1::CAMTA1/YAP1::TFE3* fusion proteins, which – although valuable for the investigation of oncogenic signaling – may not fully reproduce the broader molecular context of an established patient tumour, including additional alterations that may influence therapeutic response [12–17]. *In vivo* xenograft studies have also been limited, with few models demonstrating consistent tumour growth, long-term tolerability, or comprehensive histopathological validation. Furthermore, many reported systems focus primarily on mechanistic biology rather than establishing reproducible therapeutic evaluation platforms [18].

To bridge this translational gap, we optimized and characterized a subcutaneous (SC) cell line-derived xenograft (CDX) model of EHE using a previously established patient-derived xenograft (PDX)-derived human EHE cell line in immunodeficient mice [19]. Optimization of the cell inoculation density maintained reproducible tumour engraftment while improving animal tolerability and extending the longitudinal observation window for therapeutic studies. The resulting tumours retained key histopathological features of human EHE. We further characterized tumour proliferation, cell death, necrosis, and mTOR pathway activity and evaluated the antitumour and pharmacodynamic effects of nab-sirolimus, an albumin bound nanoparticle formulation of sirolimus that is entering clinical evaluation for EHE (SARC046 [20]). This human EHE-derived xenograft model therefore complements existing engineered endothelial and murine EHE models by providing a practical reproducible system for longitudinal preclinical therapeutic evaluation.

## 2. Materials and Methods

### 2.1 Materials

Dulbecco’s Modified Eagle Medium/Nutrient Mixture F-12 (DMEM/F12) media, fetal bovine serum (FBS), penicillin-streptomycin solution, and phosphate buffered saline (PBS) (pH 7.4) were purchased from Life Technologies (Burlington, ON, Canada). Cell Counting Kit-8 (CCK-8) was purchased from GLPBIO (Montclair, CA, USA). Matrigel™ Membrane Matrix was purchased from Corning (Corning, NY, USA). Sirolimus, everolimus, and trametinib powder were purchased from Adooq Biosciences (Irvine, CA, USA). Lyophilized nab-sirolimus (FYARRO®) in its original commercial packaging was provided by Dr. Albiruni Razak’s team at the Princess Margaret Cancer Centre (Toronto, ON, Canada).

### 2.2 Cell line and authentication

The EHE cell line used in this study was obtained and has been previously described and molecularly characterized by the Molecular Pharmacology group at the Fondazione IRCCS Istituto Nazionale dei Tumori, Milan, Italy. In the original report, the presence of the canonical WWTR1 (TAZ)–CAMTA1 fusion was confirmed by fluorescence in situ hybridization (FISH) [19].

EHE cells were cultured in DMEM/F-12 complete medium supplemented with 10% FBS and 1% penicillin-streptomycin at 37°C in a humidified incubator with 5% CO₂. Cells were passaged at 70-80% confluence using trypsin-EDTA and used for *in vivo* implantation between passages 3-10.

Cell line identity short tandem repeat (STR) profiling was performed at the Centre for Applied Genomics, at The Hospital for Sick Children (Toronto, Canada). Cell cultures were screened for mycoplasma and common pathogens by Charles River Research Animal Diagnostic Services (Wilmington, MA, USA), and all results were negative prior to initiation of animal studies.

### 2.3 *In vitro* cytotoxicity

Cytotoxicity studies on the EHE cell line were conducted using the CCK-8 assay as previously described [21]. Briefly, EHE cells were seeded into 96-well plates at 6000 cells/well and allowed to adhere for 24 hours prior to treatment. Cells were then exposed to a range of candidate therapeutics at various treatment concentrations and incubated for 72 hours. After the treatment duration, culture medium was replaced with fresh medium containing 10% CCK-8 reagent and incubated for 2.5 hours at 37 °C for colour development. The resultant UV absorbance was read on a microplate reader (Cytation5, BioTek, Winooski, VT, USA) at 450 nm and used to determine cell viability relative to drug response. Half-maximal inhibitory concentrations (IC₅₀) were calculated using GraphPad Prism 10.0 (GraphPad Software, San Diego, CA, USA) by fitting a variable slope logistic dose–response model.

### 2.4 Animals and xenograft establishment

All animal studies were approved by and conducted in accordance with the guidelines of the Animal Care Committee of the University Health Network (UHN, Toronto, ON, Canada; AUP 7034).

For tumour implantation, EHE cells were harvested during log-phase growth and resuspended in a 1:1 v/v mixture of PBS and Matrigel. Cell suspensions were maintained on ice prior to subcutaneous injection into the right flank of 6–8-week-old female SCID mice (100 µL total volume) at densities of 0.25, 0.5, 1, 2.5, or 5 × 10⁶ cells per mouse.

Subsequent xenografts were followed over time with palpable tumours measured twice weekly using digital calipers. Tumour volume was calculated using the modified ellipsoid formula:

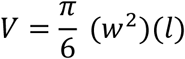

where *V* represents tumour volume, *w* represents tumour width and *l* represents tumour length.

Mice were monitored throughout the study for body weight and clinical condition (posture, grooming, mobility, and food and water intake) with humane endpoint established as an observation of either of the following conditions: tumour dimensions exceeding 15 mm in any direction, >20% body weight loss, or body condition score of less than 1.5.

At endpoint, animals were euthanized and subjected to gross pathological evaluation. Tumours were excised, fixed in 10% neutral-buffered formalin, paraffin-embedded, and processed for histopathological and immunohistochemical analyses.

### 2.5 Histology and immunohistochemistry

All histological and immunohistochemical staining was performed at the STTARR Histopathology Core Facility, University Health Network. FFPE tumour sections (4 μm) were stained with hematoxylin and eosin (H&E) for histopathological evaluation. For immunohistochemistry, sections were deparaffinized, rehydrated, subjected to heat-induced antigen retrieval, and incubated with primary antibodies against Ki-67 (clone SP6, MA5-14520; 1:100), CAMTA1 (Abcam, ab64119; 1:100), ERG (Cell Signaling Technology, #97249; 1:100), and phospho-S6 ribosomal protein (p-S6; Ser240/244; Cell Signaling Technology, #5364; 1:500). Immunoreactivity was visualized using an HRP-DAB detection system with hematoxylin counterstaining. DNA fragmentation was assessed by TUNEL following proteinase K digestion (10 μg/mL, 30 min). Sections were incubated with terminal transferase and biotin-16-dUTP (Roche) for 1 h at 37°C, followed by detection using the VECTASTAIN ABC system (Vector Laboratories) and DAB.

### 2.6 HALO quantification

Immunohistochemistry image analysis was performed using the HALO® Image Analysis Platform (Indica Labs Inc., Albuquerque, NM, USA). Briefly, a Random Forest classifier was trained on representative tissue regions to distinguish viable tumour from necrotic and non-tumour areas and was applied consistently across all samples. Staining was quantified within viable tumour regions using the HALO CytoNuclear analysis algorithm, with optimized thresholds used to classify staining intensity and determine percentage positivity and H-score.

### 2.7 *In vivo* efficacy study

#### 2.7.1 Preparation of nab-sirolimus

Nab-sirolimus solution for *in vivo* studies was prepared by dissolving commercially packed lyophilized powder at 1 mg/mL sirolimus in 0.9% saline solution, followed by filtration through a 0.22 μm PES filter into a sterile rubber-sealed glass vial. Reconstituted nab-sirolimus solution was stored on ice and used within 6 hours.

#### 2.7.2 Treatment schedule design

Treatment was initiated once tumours reached approximately 200 mm³. Mice were randomized to receive either saline control or nab-sirolimus (5 mg/kg) twice weekly intravenously on Days 1 and 4. Dosing was continued for five weeks in accordance with the standard preclinical dosing frequency described by Aadi Biosciences [22].

Tumour volume was measured twice weekly during treatment and once per week for at least 12 weeks following treatment cessation to evaluate response durability. In addition, mice were monitored throughout the study for body weight and condition with euthanasia performed upon arrival of the above stated predefined humane endpoint.

#### 2.7.3 Pharmacodynamic study

EHE xenografts with tumour volumes of approximately 200 mm³ were randomized to receive either saline or nab-sirolimus (FYARRO®). Animals received two intravenous treatments over one week using the same dosing regimen as the efficacy study. Mice were euthanized 24 hours after the second treatment, and tumours were collected for p-S6 immunohistochemical analysis.

### 2.8 Statistical analysis

Statistical analyses were performed using GraphPad Prism 10.0 (GraphPad Software, San Diego, CA, USA). Changes in tumour volume and body weight over time were analyzed using a mixed-effects model fitted by restricted maximum likelihood, with treatment, time, and treatment × time interaction as fixed effects and repeated measurements within each mouse accounted for. The overall treatment effect on tumour volume was assessed using the main effect of treatment. The p-S6 H-score and percentage of positively stained cells were compared between saline- and nab-sirolimus-treated groups using unpaired two-tailed Welch’s t-tests. P-values <0.05 were considered statistically significant.

## 3. Results

### 3.1 EHE cells exhibit sensitivity to mTOR inhibition *in vitro*

The *in vitro* cytotoxicity of the mTOR inhibitors sirolimus and everolimus together with the MEK inhibitor trametinib was evaluated in the established PDX-derived human EHE cell line. As shown in Fig. 1, treatment with all compounds resulted in a dose-dependent reduction in cell viability, with sirolimus showing a low nanomolar IC₅₀ of approximately 6.3 nM. These results confirm the previously reported sensitivity of this EHE cell line to mTOR-targeted therapy with sirolimus emerging as the promising lead candidate for further *in vivo* testing.

**Fig. 1.**
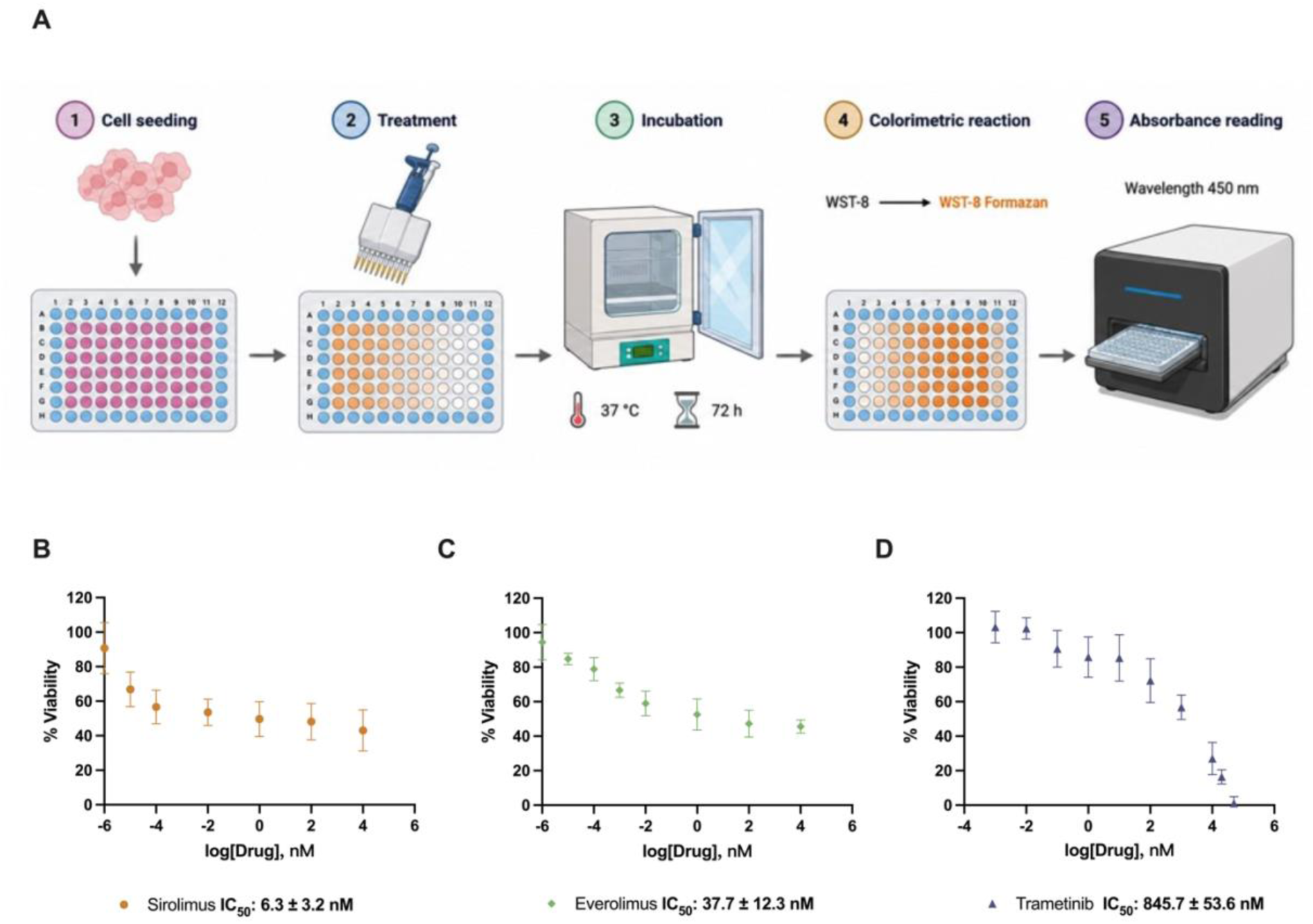
In vitro sensitivity of the PDX-derived EHE cell line to sirolimus, everolimus, and trametinib. A) Schematic of the CCK-8 Assay performed. EHE cells were treated with increasing concentrations of B) sirolimus, C) everolimus, or D) trametinib for 72 h, and cell viability was assessed using the CCK-8 assay. Cell viability was normalized to the corresponding vehicle-treated control. Sirolimus and everolimus produced lower IC₅₀ values than trametinib, indicating greater sensitivity of the EHE cell line to mTOR inhibition over the tested concentration ranges. Data are presented as mean ± SD from at least three independent experiments. IC₅₀ values are presented as mean ± SD.

### 3.2 Optimization and Characterization of a Subcutaneous EHE Xenograft Model

To enable *in vivo* evaluation of sirolimus-based therapies, a SC, CDX model of EHE was established in immunodeficient SCID mice. Initial studies comparing inoculation densities of 2.5 × 10⁶ and 5 × 10⁶ cells resulted in high take rates (i.e., 100%) and rapid tumour formation, with palpable tumours observed within 7 days. However, despite early establishment, tumour growth plateaued at approximately 200 mm³ by week 7 for both treatment densities, yielding no significant differences in tumour growth kinetics between the tested groups (Fig. 2A). This observation was accompanied by progressive body weight loss beginning around week 3 which resulted in early study discontinuation following the achievement of humane endpoints (Fig. 2B).

**Fig. 2.**
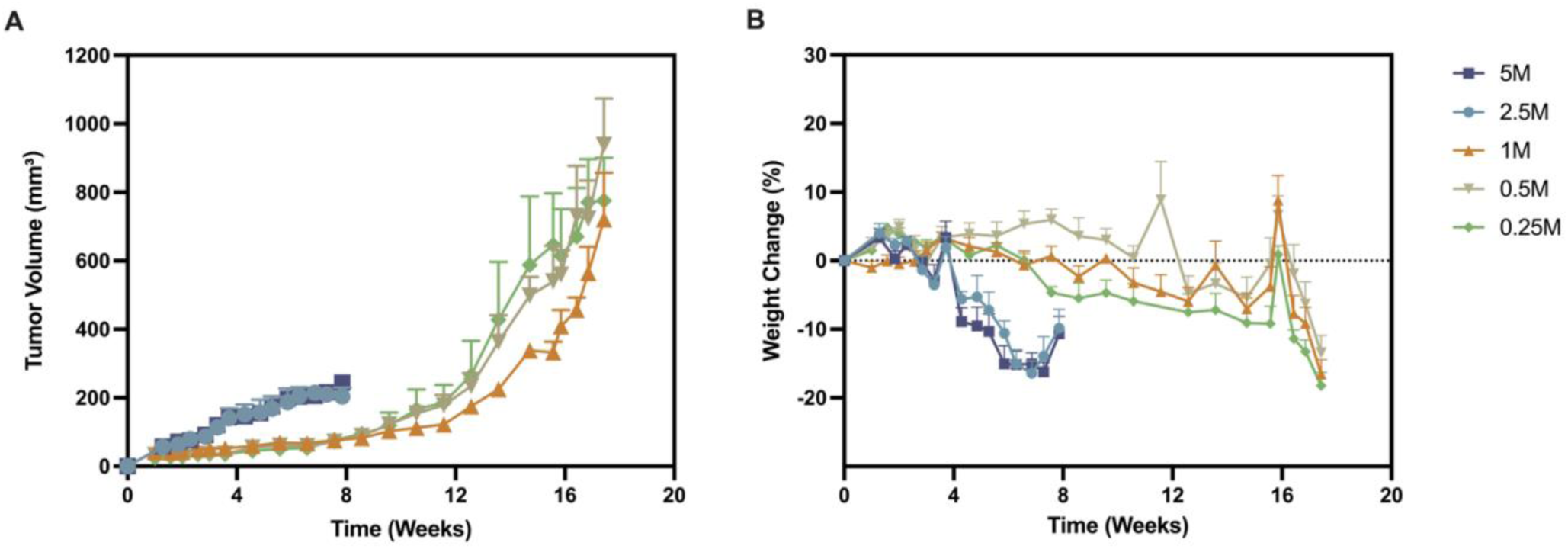
Optimization of EHE cell inoculation density for establishment of a subcutaneous xenograft model. Female SCID mice were inoculated subcutaneously with 0.25, 0.5, 1, 2.5, or 5 × 10⁶ EHE cells. A) Tumour growth and B) percent body-weight change was monitored over time. Lower cell inoculation densities resulted in slower tumour growth, allowing selection of an appropriate inoculation density for subsequent therapeutic studies. Data are presented as mean ± SEM (n = 3 per group). Only upper error bars are shown for clarity.

Gross necropsy of animals in both groups revealed abnormalities, including enlargement of the submandibular lymph nodes and abnormal pulmonary discoloration and morphology, which prompted further histological evaluation. Histological examination of the major organs, including the lungs, liver, lymph nodes, gastrointestinal tract, kidneys, spleen and bone, showed no evidence of EHE infiltration under the condition examined. Lung sections lacked the characteristic histomorphological features of EHE but exhibited diffuse vascular congestion and intra-alveolar accumulation of red blood cells, indicating a non-neoplastic pulmonary abnormality that may have contributed to the observed systemic decline (Fig. S1).

Informed by these findings, a second experiment was conducted using lower inoculation densities (1.0, 0.5, and 0.25 × 10⁶ cells) with the goal of reducing disease burden, preserving overall animal health, and establishing a broader therapeutic window for therapeutic evaluation studies. All subsequent groups maintained a 100% tumour take rate with tumours becoming palpable within 2 weeks and entering exponential growth between 6-8 weeks post-inoculation. In contrast to the high cell density groups, mice inoculated with lower cell densities remained clinically stable for a longer duration with tumours reaching volumes of approximately 1,000 mm³ prior to endpoint (Fig. 2). As no significant differences in tumour growth kinetics were observed across these groups, 0.25 × 10⁶ cells were selected as the optimal inoculation density.

### 3.3 Histopathological and Immunohistochemical Characterization

Histological evaluation of tumour sections by H&E staining revealed features consistent with EHE disease pathology, including abundant myxohyaline stroma, focal vacuolization, and characteristic tumour architecture (Fig. 3A). Immunohistochemistry staining further revealed strong nuclear expression of CAMTA1 and ERG throughout the tumour tissue, as consistent with EHE pathology (Fig. 3C).

**Fig. 3.**
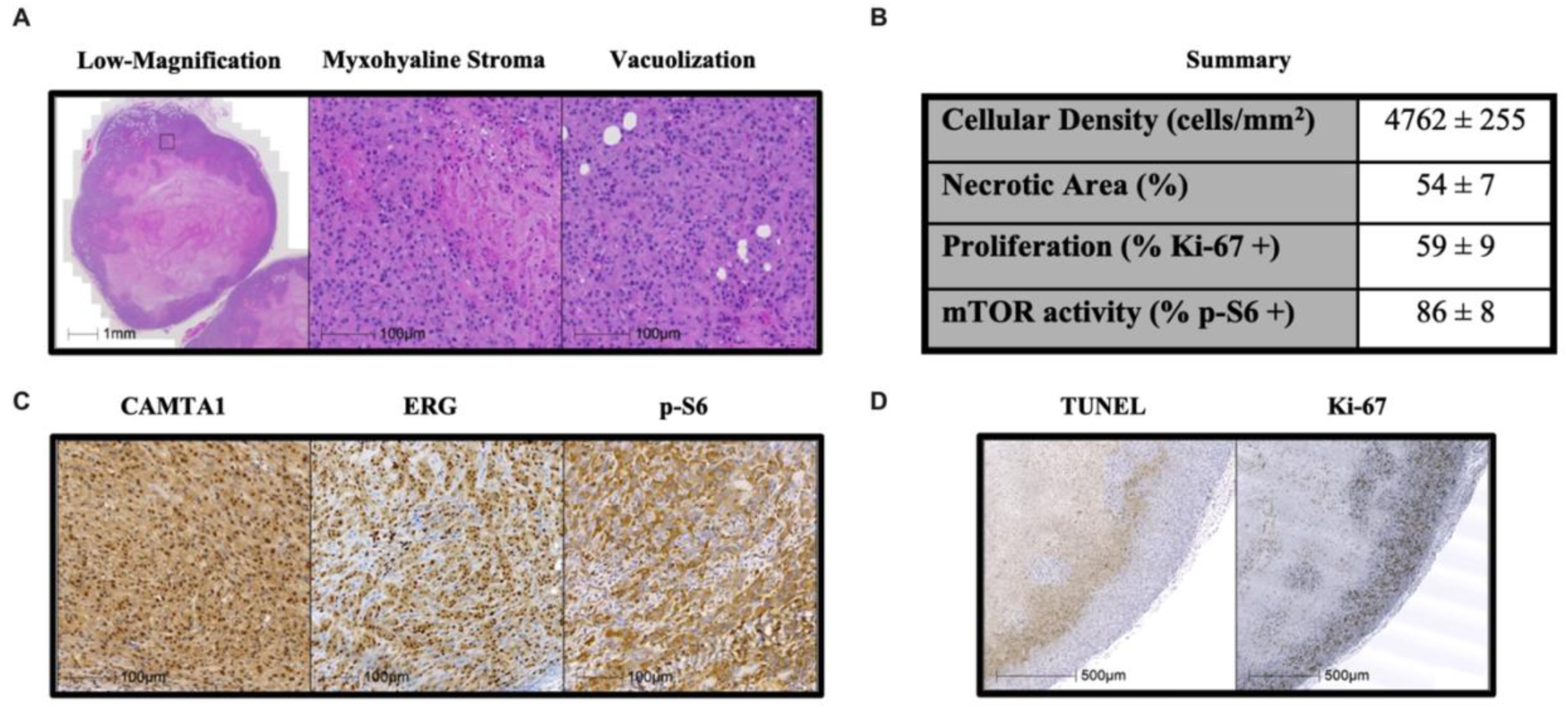
EHE xenografts retain characteristic histopathological and immunophenotypic features. A) Representative hematoxylin and eosin-stained images showing the tumour architecture, myxohyaline stroma, and cytoplasmic vacuolization. B) Quantitative summary of tumour cell density, necrotic area, Ki-67-positive cells, and p-S6-positive cells. C) Representative immunohistochemical staining for CAMTA1, ERG, and p-S6. D) Representative TUNEL and Ki-67 staining showing tumour-cell death and proliferative activity, respectively. Quantitative data are presented as mean ± SD from at least three independent tumours. Scale bars: 1 mm in the low-magnification image; 100 μm in the higher-magnification H&E and IHC images; and 500 μm in the TUNEL and Ki-67 images.

Tumour microenvironment analysis performed via H&E staining revealed a heterogeneous tumour architecture with large necrotic cores surrounded by viable EHE cells (Fig. 3A). Quantitative image analysis using HALO showed that the necrotic regions account for 54 ± 7% of tumour area, while viable regions exhibited a cellular density of 4762 ± 255 cells/mm².

Within the viable tumour regions, a significant portion of the tumour exhibited active proliferation, with 59 ± 9% of cells positive for Ki-67 (Fig. 3B). TUNEL staining was enriched at the interface between viable and necrotic regions, consistent with cell death occurring near necrotic areas (Fig. 3D). Importantly, immunohistochemical staining of p-S6, a downstream marker of mTOR pathway, was detected in 86 ± 8% of viable tumour cells (Fig. 3B, C).

### 3.4 Treatment efficacy with mTOR inhibitor

To evaluate the therapeutic responsiveness of the model to mTOR inhibition *in vivo*, tumour-bearing mice were treated with nab-sirolimus (5 mg/kg), an albumin-bound sirolimus nanoparticle approved for the treatment of advanced or metastatic PEComa. As shown in Fig. 4A, treatment administered twice weekly for five consecutive weeks resulted in significant tumour growth inhibition compared to saline controls.

**Fig. 4.**
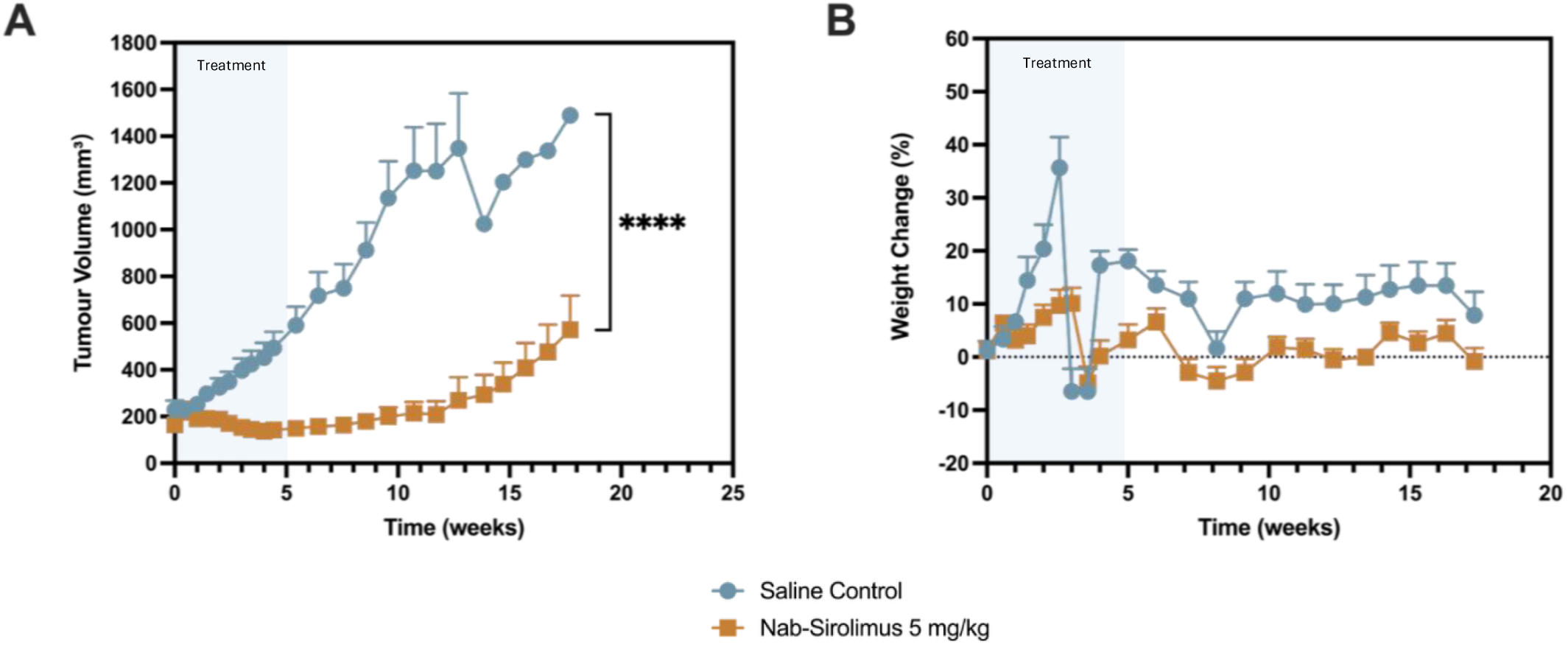
Antitumour efficacy of nab-sirolimus in mice bearing established EHE xenografts. SCID mice bearing subcutaneous EHE xenografts were randomized when tumours reached approximately 200 mm³, receiving either saline or nab-sirolimus at 5 mg/kg twice per week for five weeks. A) Tumour growth and B) percent body weight change since the first treatment day were monitored throughout the study. Nab-sirolimus treatment significantly suppressed EHE xenograft growth compared with saline treatment without sustained body-weight loss. Data are presented as mean ± SEM (n = 5 mice per group); only upper error bars are shown for clarity. ****P < 0.0001 for the overall treatment effect on tumour volume, assessed using a mixed-effects model fitted by restricted maximum likelihood.

Nab-sirolimus treatment was also generally well tolerated, with no significant or sustained reduction in body weight or body conditioning compared with saline controls (Fig. 4B).

### 3.5 Pharmacodynamic analysis of mTOR inhibition

To assess treatment-associated modulation of mTORC1 signalling, tumour p-S6 staining was evaluated by immunohistochemistry. Tumour-bearing mice treated with nab-sirolimus were shown to have reduced p-S6 positive staining compared with untreated controls (Fig. 5A). HALO quantification demonstrated a decrease in the proportion of p-S6-positive cells alongside a marked reduction in staining intensity from moderate and strong positivity toward weak positivity (Fig. 5BC). Treatment significantly reduced tumour p-S6 H-scores, consistent with suppression of downstream mTORC1 signalling (Fig. 5B).

**Fig. 5.**
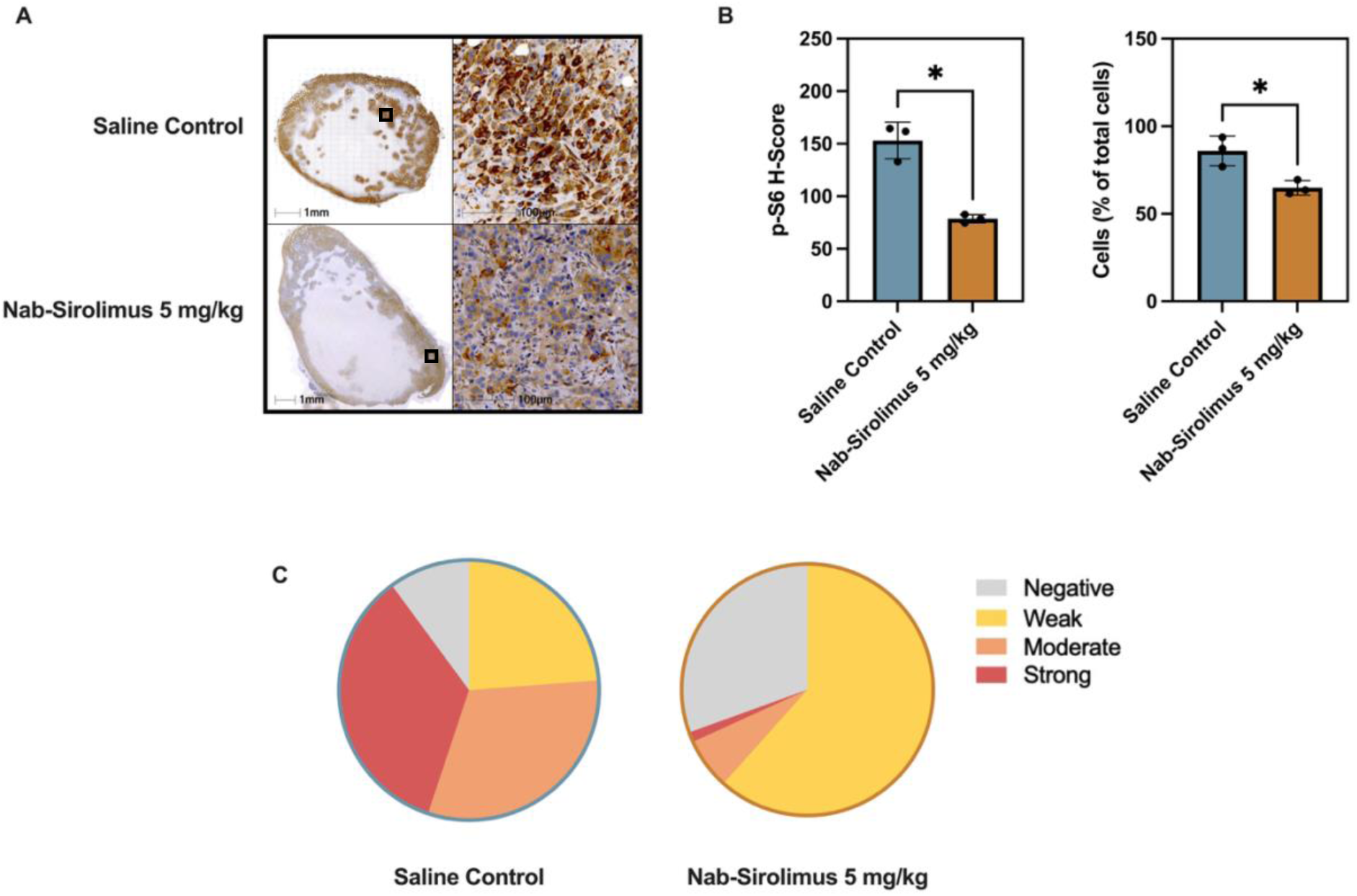
Nab-sirolimus reduces p-S6 staining in EHE xenografts. Tumours from mice treated with saline or nab-sirolimus at 5 mg/kg were assessed by p-S6 immunohistochemistry. A) Representative whole-tumour and high-magnification images of p-S6 staining. Scale bars, 1 mm and 100 μm, respectively. B) Quantification of the p-S6 H-score and percentage of p-S6-positive tumour cells using HALO. Each symbol represents an individual tumour, and bars represent mean ± SD (n = 3 per group). C) Distribution of tumour cells classified as negative, weakly positive, moderately positive, or strongly positive according to p-S6 staining intensity. * Denotes a statistically significant difference between nab-sirolimus treated group versus the saline control (*P* < 0.05).

## 4. Discussion

### 4.1 Optimizing cell inoculation density extends the therapeutic evaluation window while retaining EHE diagnostic markers

Optimization of the SC tumour engraftment protocol revealed that inoculation with high cell densities was associated with rapid declines in body condition despite slow and plateauing growth of the primary tumour. Although gross necropsy findings revealed abnormal pulmonary colour and morphology, raising concern for possible systemic involvement, histological evaluation of major organs did not reveal evidence of EHE infiltration. Instead, pulmonary tissues showed vascular congestion and intra-alveolar red blood cell accumulation, suggesting a non-neoplastic pulmonary condition. Although the underlying mechanism remains unclear, these findings may reflect a systemic inflammatory or vascular response associated with excessive tumour cell burden.

Notably, inoculation with high cell densities, including 2.5 and 5 million cells, produced SC tumours with limited growth duration and non-sigmoidal growth profiles prior to rapid host decline, making the model unsuitable for longitudinal preclinical drug testing. This observation is consistent with the pilot findings reported by Stacchiotti et al., whose 5-million-cell SCID mouse model confirmed the in vivo tumour-forming capacity of the EHE cell line but was similarly limited to a short growth duration before study termination.

Based on these parallel findings and our host body condition observations, we hypothesized that reducing the initial cell inoculant density could improve tolerability while preserving tumour engraftment. Consistent with this hypothesis, reducing the initial cell inoculant density significantly improved host tolerability and extended the observation period. Although injection of lower cell quantities lengthened the tumour latency period, the 100% take rate was maintained, with tumours exhibiting classical sigmoidal growth kinetics. This extended the therapeutic evaluation window, allowing treatment effects to be monitored over a more clinically meaningful duration.

Pathological assessment demonstrated that xenografts generated using the optimized protocol retained nuclear CAMTA1 and ERG expression, supporting preservation of the characteristic immunophenotype of conventional WWTR1–CAMTA1-positive EHE and confirming endothelial differentiation [23,24]. Together, these findings demonstrate that the optimized CDX retains key diagnostic features of human EHE while providing a growth profile suitable for preclinical drug testing.

### 4.2 EHE tumours exhibit core necrosis and baseline p-S6 expression

Histological and quantitative image analyses of untreated tumours revealed a necrotic core surrounded by a rim of viable tumour cells, consistent with the histological observations previously reported [19]. TUNEL-positive cells were predominantly observed at the interface between viable and necrotic tumour regions, consistent with increased apoptotic activity in this transitional zone. Within the viable regions, Ki-67 staining demonstrated high proliferative activity, with 59 ± 9% of viable cells staining positive. Together, these findings support a highly proliferative tumour phenotype and are consistent with the histopathological characteristics reported for the original patient-derived tumour [19].

To characterize baseline mTORC1-associated signalling in the CDX and establish a reference for subsequent pharmacodynamic assessment, resected tumours were additionally stained for p-S6 (Ser240/244), a downstream marker of mTORC1 activity [25,26]. As shown in Fig. 3, 86 ± 8% of viable EHE cells were positive for p-S6, indicating widespread basal p-S6 expression and supporting the use of this model for evaluating mTOR-targeted therapies. This finding provided a useful baseline for assessing treatment-associated changes in p-S6 *in vivo*.

### 4.3 Nab-sirolimus suppresses EHE xenograft growth and reduces tumour p-S6 staining

An efficacy study was conducted to determine whether the optimized model could support longitudinal therapeutic evaluation using a clinically relevant mTOR-targeted treatment. Sirolimus was selected based on its established activity in EHE, with clinical studies demonstrating prolonged disease stabilization in patients with progressive advanced EHE, particularly in the absence of serosal effusions [11]. Consistent with these clinical findings, sirolimus has also demonstrated antitumour activity in the corresponding EHE PDX model and its paired cell line [19]. Our *in vitro* studies further confirmed sensitivity of the EHE cells to mTOR inhibition. For *in vivo* therapeutic validation, nab-sirolimus was selected as an intravenously administered, albumin-bound formulation of sirolimus with established clinical activity and regulatory approval in malignant PEComa [27,28]. Importantly, nab-sirolimus is currently being evaluated in a Phase II clinical trial in patients with progressing or symptomatic EHE [20], further supporting its translational relevance to this disease.

Treatment with nab-sirolimus at 5 mg/kg delayed tumour progression relative to saline control and was well tolerated, with no significant body-weight loss observed during treatment. The response to systemic mTOR inhibition was consistent with that previously reported in the corresponding PDX model [19]. Although tumour regrowth was observed six weeks after the final treatment, nab-sirolimus produced a clear and measurable period of tumour-growth suppression.

The initial tumour regression followed by subsequent regrowth suggests that mTOR inhibition suppressed tumour growth without completely eliminating viable tumour cells. This pattern is consistent with clinical observations in EHE, where mTOR inhibitors are more commonly associated with prolonged disease control rather than complete tumour eradication [11,29].

Pharmacodynamic analysis further provided evidence of treatment-associated modulation of p-S6. Following one week of treatment, corresponding to two doses of nab-sirolimus at 5 mg/kg, treated tumours exhibited significantly reduced p-S6 staining compared to saline controls. The reduction in p-S6 positivity was consistent with suppression of mTORC1 signalling. Together, the tumour-growth and pharmacodynamic findings demonstrate that the CDX model can support both longitudinal therapeutic efficacy assessment and evaluation of downstream mTORC1 pathway modulation *in vivo*.

### 4.4 Complementary value of the EHE CDX for therapeutic evaluation

Historically, engineered cellular and mouse models have been instrumental in establishing the oncogenic role of WWTR1(TAZ)–CAMTA1 and advancing our understanding of EHE biology. Expression of the fusion in non-EHE cellular backgrounds, including HEK293 and NIH3T3 cells, has enabled controlled investigation of its transforming and transcriptional functions, while genetically engineered mouse models have demonstrated that endothelial expression of WWTR1–CAMTA1 is sufficient to generate tumours that recapitulate key features of human EHE [12–17].

Although engineered cellular and mouse models have been invaluable for studying WWTR1(TAZ)–CAMTA1-driven tumour biology, their reliance on experimentally introduced or enforced fusion expression may not fully capture the broader molecular context of an established patient tumour, including co-existing and adaptive alterations that can influence therapeutic response [30,31]. This consideration does not diminish their value for mechanistic or therapeutic studies but highlights the complementary role of patient EHE-originating models in translational drug evaluation.

Recently, Stacchiotti et al. established a *WWTR1::CAMTA1*-positive human EHE PDX and a corresponding PDX-derived human EHE cell line, and demonstrated the tumourigenicity of this cell line following xenotransplantation [19]. While these studies provided an important foundation for developing patient-derived EHE models, the initial xenotransplantation conditions were associated with a limited tumour-growth duration and small maximal tumour volumes prior to study termination. Building on this work, the present study optimized the xenotransplantation conditions to improve host tolerability, extend the tumour-growth window, and facilitate longitudinal therapeutic evaluation. Under the optimized conditions, the resulting tumours retained key diagnostic features of EHE and showed a measurable response to mTOR inhibition. This optimized CDX therefore provides a practical and accessible complement to existing EHE models for preclinical therapeutic studies, particularly when access to PDX models is limited.

### 4.5 Limitations

Several limitations of this model should be acknowledged. First, SC engraftment does not fully reproduce the native tissue microenvironments in which EHE arises. Although this approach facilitates longitudinal tumour monitoring and therapeutic assessment, site-specific interactions between EHE cells and their surrounding microenvironment may not be fully captured.

Second, the use of immunodeficient SCID mice limits evaluation of adaptive immune responses and immunotherapeutic strategies [32]. Immunocompetent genetically engineered or syngeneic models may therefore be better suited for investigating immune-tumour interactions and therapies that target the immune system.

Third, the model was generated from a single *WWTR1::CAMTA1*-positive cell line and therefore does not capture the full molecular and clinical heterogeneity of EHE, including *YAP1::TFE3*-positive disease.

Finally, pharmacodynamic assessment of mTOR pathway modulation was limited to p-S6 immunohistochemistry. Although reduced p-S6 staining following nab-sirolimus treatment was consistent with suppression of downstream mTORC1 signalling, additional pathway markers (e.g. 4EBP1) and orthogonal validation approaches, such as Western blotting, would strengthen mechanistic interpretation.

## 5. Conclusion

In conclusion, we optimized and comprehensively characterized the SC xenotransplantation of a previously established PDX-derived human EHE cell line for longitudinal therapeutic studies. By reducing the cell inoculation density, reliable tumour engraftment was maintained while host tolerability and the tumour-growth window were substantially improved, overcoming key practical limitations of the initial xenotransplantation conditions. The resulting xenografts retained key diagnostic features of EHE and exhibited measurable basal p-S6 expression. Under these optimized conditions, nab-sirolimus treatment produced a clear antitumour response accompanied by reduced p-S6 staining, demonstrating that the model can support both longitudinal efficacy assessment and pharmacodynamic evaluation.

Given the scarcity of experimentally accessible models for this ultra-rare vascular sarcoma, these optimized conditions address an important preclinical gap by providing a practical and reproducible *in vivo* system for evaluating candidate therapies. While complementary to existing engineered and patient-derived models, this CDX offers a readily deployable approach for therapeutic studies in EHE and may facilitate broader preclinical investigation of new treatment strategies.

## Disclosure statement

No potential conflict of interest was reported by the authors.

## Data availability statement

The data supporting the findings of this study are available from the corresponding author upon reasonable request.

## Declaration of generative AI use

During preparation of this manuscript, the authors used ChatGPT to assist with language editing and improve clarity. The authors reviewed and revised the output and take full responsibility for the final manuscript.

## Acknowledgements

These studies were supported by the Sarcoma Cancer Foundation of Canada through the Multi-pronged Canadian Research in Epithelioid Haemangioendothelioma (PRO CARE EHE) project. Zilu Chen holds a Centre for Pharmaceutical Oncology (CPO) scholarship. The authors acknowledge the use of equipment at the CPO (University of Toronto), and the STTARR Innovation Centre (University Health Network). The generation of the EHE cell line used in this study was supported by Associazione Italiana per la Ricerca sul Cancro (AIRC; IG 29477, S. Stacchiotti), EHE Rare Cancer Charity, United Kingdom, and the EHE Foundation, United States (N. Zaffaroni and S. Stacchiotti).

## Author Contributions

**Zilu Chen:** Conceptualization, Methodology, Investigation, Formal analysis, Data curation, Visualization, Writing – original draft, Writing – review & editing. **Lucy Wang:** Conceptualization, Methodology, Project administration, Writing – review & editing. **Valentina Zuco:** Methodology, Resources. **Sandro Pasquali:** Resources, Writing – review & editing. **Silvia Stacchiotti:** Resources. **Nadia Zaffaroni:** Resources, Project administration. **Albiruni R. Abdul Razak:** Conceptualization, Resources, Methodology, Supervision, Project administration, Funding acquisition. **Christine Allen:** Conceptualization, Supervision, Project administration, Funding acquisition, Writing – review & editing.

## Notes

### Competing Interest Statement

The authors have declared no competing interest.

