## Supplemental Figures for "Optimization of a Subcutaneous Epithelioid Hemangioendothelioma Xenograft Model for Evaluation of Preclinical Therapeutics"

### Supplementary Information

#### EHE Cell Line STR

| Sample File | Sample Name | Panel | Marker | Allele 1 | Allele 2 |
|---|---|---|---|---|---|
| GP10-WAN30823 | EHE_1_Cell_Line | GenePrint_10 | AMEL | X | X |
| GP10-WAN30823 | EHE_1_Cell_Line | GenePrint_10 | CSF1PO | 10 | 12 |
| GP10-WAN30823 | EHE_1_Cell_Line | GenePrint_10 | D13S317 | 11 | 11 |
| GP10-WAN30823 | EHE_1_Cell_Line | GenePrint_10 | D16S539 | 9 | 12 |
| GP10-WAN30823 | EHE_1_Cell_Line | GenePrint_10 | D21S11 | 28 | 30.2 |
| GP10-WAN30823 | EHE_1_Cell_Line | GenePrint_10 | D5S818 | 9 | 11 |
| GP10-WAN30823 | EHE_1_Cell_Line | GenePrint_10 | D7S820 | 8 | 12 |
| GP10-WAN30823 | EHE_1_Cell_Line | GenePrint_10 | TH01 | 6 | 6 |
| GP10-WAN30823 | EHE_1_Cell_Line | GenePrint_10 | TPOX | 8 | 8 |
| GP10-WAN30823 | EHE_1_Cell_Line | GenePrint_10 | vWA | 16 | 16 |

Human-specific Ku80 immunohistochemistry was used to assess potential EHE infiltration in the major organs. EHE xenograft tissue and healthy mouse tissue served as positive and negative tissue controls, respectively. Representative lung sections from EHE xenograft-bearing mice showed no detectable Ku80-positive human cells, while corresponding hematoxylin and eosin staining showed no histological evidence of EHE infiltration or metastasis.

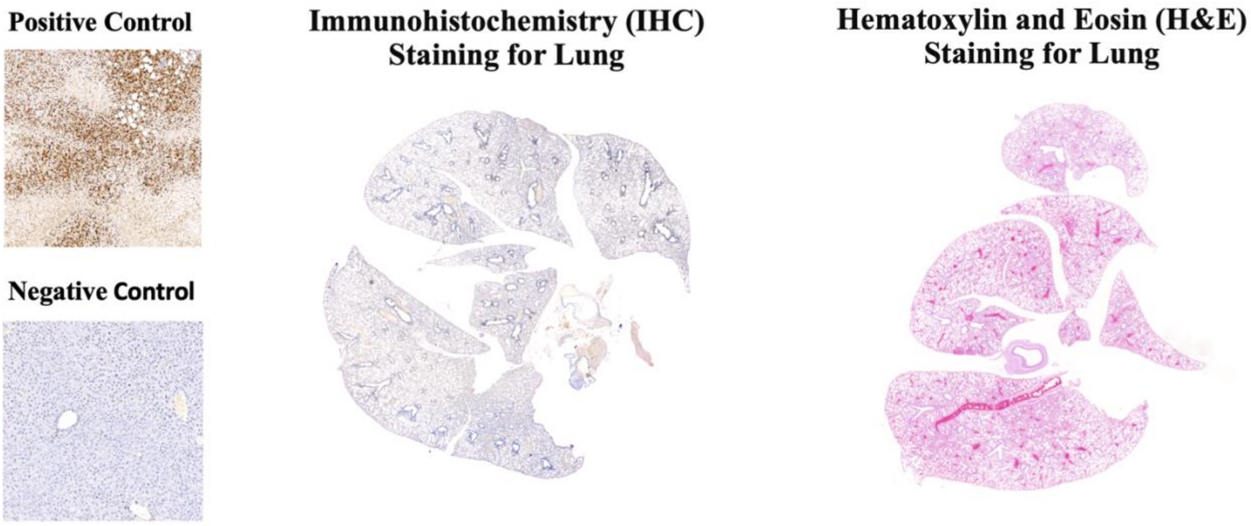


**Supplementary Fig. 1**. Histological evaluation of the lung revealed no evidence of human EHE cell infiltration.
